# Precisely phase-locked acoustic stimuli globally enhance slow oscillations, but depress fast spindles

**DOI:** 10.1101/2025.03.04.639851

**Authors:** Dominik P. Koller, Kim Hubregtse, Anna C. van der Heijden, Lucia M. Talamini

## Abstract

**Introduction:** Several studies have shown manipulation of slow oscillations (SO) and sigma power through auditory stimulation during sleep. Most of the evidence, however, regards effects immediately following stimulation rather than enduring effects. Moreover, effects on discrete fast and slow spindles have as yet not been assessed.

**Materials and Methods:** Here we use a modeling-based approach to predict upcoming oscillatory activity in the EEG and precisely phase-lock subtle acoustic stimuli to the start of the SO positive deflection. We assess the effects of stimulation on discrete slow oscillations, fast and slow spindles in the seconds after stimulation and on the longer term. We relate our findings to observations at the level of spectral measures and stimulus evoked responses.

**Results:** Our observations show that slow wave measures were consistently increased, as apparent in measures of discrete SO’s, SO and delta power and deflections in the ERP. On the other hand, fast spindle measures showed a temporally localized increase during a stimulus-induced SO positive deflection around 1 second after stimulation, but were globally decreased, both on the short and long term. The latter was apparent in measures of discrete fast spindles and PSD across longer periods of sleep.

**Conclusions:** Acoustic stimuli, precisely phase locked to the SO onset, increase SO’s and delta power globally, therewith deepening sleep. On the other hand, fast sleep spindles are globally depressed. This appears to be due, in part, to interruption of ongoing spindles by the stimulus and may furthermore reflect a depressing influence of slow oscillations on fast spindle dynamics. Future studies could evaluate the therapeutic potential of sleep deepening with acoustic stimulation in clinical populations who suffer from reduced deep sleep, such as in insomnia or post-traumatic stress disorder.

## Introduction

Attempts to non-invasively enhance or manipulate sleep oscillations have been ongoing for over a decade. Especially slow oscillations (0.5 to 1.5 Hz) and sleep spindles (11 - 16 Hz), the hallmarks of non-rapid eye movement (NREM) sleep, have been the subject of several studies. These investigations provide evidence that rocking [1,2], transcranial magnetic stimulation [3], transcranial alternating current stimulation [4], intracranial stimulation and auditory stimulation [5–9] can influence sleep oscillations and, in some cases, improve memory consolidation.

Most recent experiments have used auditory stimulation to modulate slow oscillations and sleep spindles [10–14]. In view of their non-invasive nature and practical convenience, auditory stimulation methods bear promise for clinical and home-use applications. Additionally, the rich content-conveying properties of auditory cues facilitate further investigation into the role of sleep waves in memory processes. Tracking ongoing brain activity and aligning auditory stimuli with slow oscillations through closed-loop procedures, has turned out to be particularly effective in manipulating slow oscillations and spindles [5–9,15–17]. Furthermore, the precise phase at which slow oscillations are targeted importantly determines the response. In particular, it has been shown that sounds targeted to the start of the slow oscillation up-wave, compared to down-wave targeted sounds, produce a more pronounced subsequent up-wave, coupled to enhanced spindle and beta activity [6,18].

Studies using such approaches have mainly focused on the analysis of acute responses to auditory stimulation, up to a few seconds following a stimulus, or during rhythmic stimulation around the slow oscillation frequency (∼0.5-1.5 Hz). A few studies that assessed longer-term effects (i.e. more than a few seconds away from stimulation) reported no notable changes to sleep architecture [6–9,15]. Of note, in some studies using frequent, rhythmic stimulation, power increases around the stimulation frequency were observed [9,19,20]. Again, these observations likely reflect the immediate effects of a rhythmic input on the EEG signal. Studies assessing the effects of rhythmic stimulation in more detail have, however, shown that the EEG response to subsequent pulses in an acoustic stimulus train quickly fades out [5,8,21].

In view of the above, the first aim of this study was to assess whether single acoustic stimuli, precisely phase-targeted to SO up-waves, would lead to more enduring effects on oscillatory sleep dynamics. These could translate to global, sleep-enhancing effects with potentially important consequences for sleep quality, restorative properties, and information processing. The use of single sounds ensures that every stimulus can be precisely phase targeted. Conversely, with rhythmic stimulation, applied in most studies thus far, untargeted stimuli might fail to enhance, or even disturb ongoing oscillatory activity [22].

As indicated above, studies thus far have shown that acoustic stimulation during NREM sleep may evoke a K-compex or amplified SO-like response immediately after stimulation. The induced up-wave, tends to be coupled to enhanced spindle and beta activity. As an ulterior, complementary objective, we assessed whether pulses of white noise featuring an amplitude modulation in the spindle frequency range might be particularly effective for modulating sleep spindles.

To achieve accurate and precise phase targeting, this study uses a novel closed-loop stimulation approach, based on real-time modeling and prediction of oscillatory brain dynamics [23]. This method copes flexibly with the high time-variance of the EEG signal. We used it to aim single, white noise pulses at the transition from the negative to the positive deflection of slow oscillations, and to compare stretches of stimulated and unstimulated sleep in a within-subject, nap design.

In accordance with our main aim, we systematically analyze both short-term and longer-term responses of slow oscillations and sleep spindles to the acoustic stimulation. While slow wave characteristics, such as amplitude, duration and slope, have gotten reasonable attention in previous studies [8,9,16,24,25], sleep spindle characteristics were explored less thoroughly. Therefore, our objective was to prioritize the analysis of discrete spindle events, in addition to slow wave events. Finally, we performed analyses to understand how acoustic stimulation induced changes at the level of discrete slow wave and spindle events relate to spectral EEG measures and acoustic stimulus evoked responses.

Accumulating evidence suggests that the traditional spindle band (11-16 Hz) hosts two types of sleep spindles, slow spindles (11-13 Hz) and fast spindles (13-16 Hz) [26,27]. Notably, slow and fast spindles have different hemodynamic responses [28], topographic distributions [29], circadian modulation [30,31], hereditability [27], and occur at different phases of slow oscillations [32], likely reflecting distinct generating mechanisms and functional roles. Therefore, we conducted our analyses separately for slow and fast spindles.

## Methods

### Participants

Twenty-eight subjects (20.36 ± 1.55 yr., 13 males) from the Faculty of Social and Behavioral Sciences of the University of Amsterdam, with no prior history of sleep disorders or other neurological, psychological, or psychiatric disorders, participated in the experiment for university credit points. We excluded 14 datasets of the initial sample because the subjects could not fall asleep (2), did not reach consolidated sleep stage N3 (3) or did not go through at least five experimental blocks in each condition (9). Thus, 14 subjects (20.59 ± 1.72 yr., 8 male) were included in our analyses. The participants abstained from alcohol, caffeine-containing drinks and any other sedative or stimulating substances for at least 24h prior to the investigation. They were asked to wake up three hours before their usual wake-up time, but at 6:00 AM at the latest, on the day of the experiment. All volunteers gave written informed consent before participating. Our study was approved by the ethical committee of the University of Amsterdam.

### Auditory Stimuli

Two different auditory stimuli were generated with Audacity® software version 2.2.2 (Audacity Team, 2017): a pure white noise stimulus and a white noise stimulus amplitude-modulated by a 14 Hz sine wave between 0 and 100% of its original amplitude. The amplitude-modulated stimulus was created with the specific goal of inducing sleep spindles by using a frequency within the spindle range. Both stimuli had a duration of 286 ms, equivalent to four cycles at the modulation frequency. The stimuli were presented with two stereo speakers (GigaWorks T20 Series II, Creative Technology Ltd., Jurong East, Singapore) placed approximately 50 cm from the head of the subject. We used 10 s loops of each sound to calibrate the speakers to an equivalent continuous sound pressure level of 42.5 dBA, as estimated with a sound level meter (NA-27, Rion Co., Ltd., Kokubunji, Tokyo, Japan).

### Procedure

On the day of the investigation, participants arrived at the sleep laboratory at 12:30 PM. Following completion of the screening questionnaires, they were fitted with 62 EEG electrodes according to the international 10-10 system (64-channel WaveGuard cap, ANT, Enschede, The Netherlands), two electrodes placed on the mastoids, bipolar horizontal EOG electrodes placed next to the canthus of the left and right eye, bipolar vertical EOG electrodes above and below the right eye, and submental bipolar EMG electrodes. Signals were sampled at 512 Hz using 72-channel Refa DC amplifiers and an in-house adapted version of Polybench software (TMSi, Oldenzaal, The Netherlands).

After electrode placement participants got a 3h sleep opportunity. Lights were switched off between 13:24 and 14:17 (range across all participants). Upon visual detection of N3 sleep according to standard American Academy of Sleep Medicine (AASM) criteria [33], the phase-tracking and stimulation algorithm was started. Stimulus presentation mode was temporarily switched off if the participants showed signs of arousal or shifts to any other sleep stage.

In stimulus presentation mode, the algorithm alternated between stimulation blocks (SB) and buffer blocks (BB), each lasting 30 s (Figure 1). Every SB was randomly assigned to one of three stimulation modes: white noise (WN), amplitude-modulated white noise (AMWN), or sham (SHAM; EEG event marker, but no sound). Depending on the stimulation mode, a single stimulus (or event marker in SHAM) was presented to coincide with predicted zero-phases in ongoing slow oscillations; that is, the transition from the negative to the positive deflection of a slow wave. After each single-sound stimulation the algorithm was halted for 4.5 s to ensure stimulation-free periods in which we assessed short-term (0 to 3 s after stimulus onset) effects of stimulation. The BBs are stimulation-free periods that allow wash out of longer-lasting effects of the auditory stimulations occurring during SBs.

**Figure 1.**
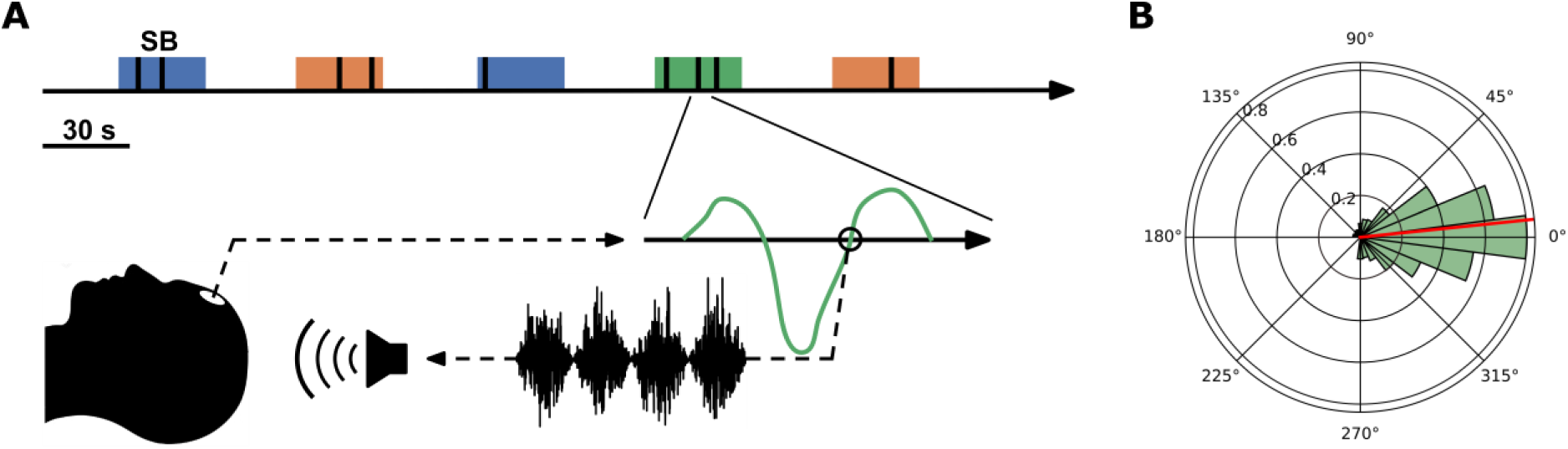
Experimental design and phase-targeting accuracy. (A) The algorithm alternates between 30 s stimulation blocks (SB), during which WN, AMWN, or SHAM stimulate (black bars) are released (each condition is indicated with a different color) and buffer blocks (BB; periods between SBs). (B) The algorithm is tracking the phase of ongoing slow wave activity in channel Fpz, while targeting stimuli to slow waves’ negative-to-positive transition (indicated for one stimulus in an AMWN block). At the transition, a sound (indicated by the amplitude-modulated waveform) is played via speakers to the participant. (B) The distribution of stimuli across the slow wave phase is plotted in a circular histogram. The red line indicates the circular mean of 5.92 ± 52.83 deg. The negative-to-positive transition, positive deflection peak, positive-to-negative transition and negative deflection peak correspond to 0 deg, 90 deg, 180 deg and 270 deg respectively.

### Phase-tracking Algorithm

As high-density EEG was being recorded, a copy of the Fpz-M1 signal was batch transferred to a second computer for real-time analysis. The data was received into a buffer holding the most recent 10 seconds of data. A novel oscillatory phase prediction procedure, named modelling-based closed loop neurostimulation (M-CLNS), developed in house (collaboration with Okazolab Ltd, Delft, The Netherlands), performs non-linear sine fitting on the most recent one-second of unfiltered EEG signal and, at each iteration, checks the following criteria for stimulus release: (1) the fitted sine is in a predefined frequency range of interest (here 0.5-1.5 Hz), (2) the fitting error, in terms of the mean standard error across the fitting segment, is below threshold (0.3) (3) the predicted target phase occurs within a narrow time limit (34 ms). If all criteria are met, the sound stimulus is released to coincide with the upcoming predicted target phase. The real-time phase prediction algorithm was implemented in EventIDE software (Okazolab Ltd, Delft, The Netherlands).

### Preprocessing

The data was preprocessed with the open-source, Python-based (Python Software Foundation, Python version 3.6.5) MNE software [34]. First, data was re-referenced to the left and right mastoid electrodes and bandpass filtered between 0.3 to 40 Hz with a zero-phase finite impulse response filter, using the windowed time-domain design method (Hamming window; filter length of 11.002 s). Next, we visually inspected the signal traces and power spectral densities of all electrophysiological measurements to identify noisy channels and assess general data quality. Bad channels were repaired with a spherical spline interpolation. Two independent raters scored the recordings for sleep-wake stages based on standard criteria [33] using the open-source software Wonambi (Piantoni & O’Byrne, 2013). Interrater agreement was 81% (Cohen’s Kappa = 0.74) in line with common interrater reliability [35]. Subsequently, condition blocks (CB) from sleep stage N3 were extracted for further analyses. These CBs started with the onset of the first stimulus in a SB and were defined to last 30 s. With this definition of CBs, we ensured to only include brain activity affected by prior stimulation. Notably, this method implicitly excludes SBs with no stimulus presentation. SBs where a stimulus was cut-off by the transition to a BB were also excluded. At this stage, we verified that each participant had received at least five CBs per stimulation mode (AMWN, WN or SHAM). We also visually inspected CBs’ signal traces and power spectral densities to ensure artefact-free data. Finally, to investigate the short-term effects of auditory stimulation, epochs from -1.5 to 3.5 s relative to stimulus onset were extracted from the CBs.

### Phase-targeting accuracy

We evaluated our algorithms phase-targeting accuracy using the instantaneous phase of each participants EEG-signal filtered in the slow wave frequency range (0.5 to 1.5 Hz). Instantaneous phase was estimated by applying the Hilbert transform followed by computing the phase angle from the complex-valued analytic representation. The circular mean and standard deviation of the slow wave phase at sound onset, at Fpz, across all participants and conditions were calculated.

### Power Spectral Density

Global power changes within 0.1 to 40 Hz were investigated through power spectral density (PSD). PSDs were estimated for CBs using Welch’s method with 2-second Hamming windows and 50% overlap. Next, we averaged resulting PSDs within conditions for each participant, followed by a log-total power normalization using 10 𝑙𝑜𝑔_10_(𝑃𝑆𝐷/𝑃𝑆𝐷_𝑡𝑜𝑡𝑎𝑙_ ), where 𝑃𝑆𝐷 is the power spectrum corresponding to a specific condition of a participant and 𝑃𝑆𝐷_𝑡𝑜𝑡𝑎𝑙_ is the total power contained within 0.1 to 40 Hz across all conditions (using Simpson’s rule to approximate the integral between 0.1 to 40 Hz).

### Event-related Potentials

Event-related potentials (ERPs) were obtained by averaging across all stimulation epochs for each condition, resulting in -1 to 3-second ERPs referenced to stimulus onset. No baseline correction was applied to ERPs since they were high pass filtered during preprocessing [36].

### Time-frequency Analysis

We computed time-frequency representations (TFRs) using the complex Morlet wavelet transform of the open-source Python package MNE [34]. We determined a suitable trade-off between spectral and temporal precision for slow wave and sleep spindle wavelet center frequencies by estimating the full-width half maximum (FWHM) [37] values for a range of number of cycles. We used a center frequency of 13 Hz for spindles based on median estimates by Purcell and colleagues [27]. We found that 8.5 cycles, at this center frequency, resulted in a temporal FWHM resolution of 245 ms, sufficient to resolve spindle activity nested in positive or negative slow wave deflections, assuming a median slow wave frequency of 1 Hz. The spectral FWHM resolution of 2.55 Hz (8.5 cycles at 13 Hz) is sufficient to resolve slow and fast spindles.

To resolve slow wave frequencies, we decided to use 3 cycles at 2 Hz center frequency, resulting in 562 ms and 1.11 Hz temporal and spectral FWHM, respectively. We set our lower bound to 2 Hz to avoid estimation issues [37]. The temporal resolution is sufficient to depict slow wave power changes and the spectral resolution includes low frequency content.

After optimizing for our primary objectives, we linearly increased the cycle number with frequency from 3 cycles at 2 Hz to 12 cycles at 20 Hz. Computed TFRs ranged from -1.5 to 3.5 s relative to stimulus onset with a step size of 20 ms and were estimated from 2 Hz to 20 Hz in 0.5 Hz steps. After averaging TFRs across individual epochs for each stimulation condition, we log-power normalized them with 10 𝑙𝑜𝑔_10_(𝑃/𝑃_𝑏𝑎𝑠𝑒𝑙𝑖𝑛𝑒_), where 𝑃_𝑏𝑎𝑠𝑒𝑙𝑖𝑛𝑒_ is the average power across a -1.25 to -0.25 s baseline relative to stimulus onset. Notably, the temporal FWHM is 562 ms at 2 Hz, hence this baseline sufficiently avoids leakage of boundary effects and pre-stimulus power into post-stimulus activity, while normalizing slow wave activity. Next, we averaged TFR epochs across all participants for each stimulation condition and cropped them to -1 to 3 s relative to stimulus onset to remove boundary effects.

### Spindle Detection

Before spindle detection, we visually identified slow and fast spindle frequency peaks between 9 to 17 Hz, on channels Fz and Pz, respectively, for each participant. In four participants slow spindle peaks could not be identified and were approximated by averaging across the remaining participants. Slow and fast spindle frequency bands were each defined as a 3 Hz range centered around previously defined peak frequencies.

Next, discrete slow and fast spindles were detected with the Python package Yet Another Spindle/Slow Wave Algorithm (YASA) [38], largely based on the algorithm of Lacourse and colleagues [39]. The algorithm first computes a moving RMS, with a 300 ms window and a step size of 100 ms, from EEG data filtered within defined spindle frequency bands. Then a moving correlation between the pre-processed EEG signal and the EEG signal filtered within the spindle frequency bands is computed with a sliding window of 300 ms and a step size of 100 ms. Lastly, the Short-Term Fourier Transform is computed from consecutive two-second windows with a 200 ms overlap. Moving relative power is then calculated from the ratio of power in the spindle frequency bands and the total power from 0.3 to 40 Hz. Sleep spindle candidates are identified from these signals whenever the moving RMS crosses a threshold defined by 𝑅𝑀𝑆_𝑚𝑒𝑎𝑛_ + 1.5 𝑅𝑀𝑆_𝑆𝐷_, or if the moving correlation surpasses 0.50, or if the relative power exceeds the relative power threshold calculated based on each subject’s spindle frequency bands. Spindles were detected only if two thresholds were reached simultaneously. Furthermore, spindles shorter than 0.3 s or longer than 2.5 s were discarded. We assumed spindles occurring in different channels, within 500 ms, reflect the same spindle. In these cases, we chose the spindle with the maximum relative power for further analyses. Duration, amplitude, and frequency were calculated for each detected spindle. Additionally, we computed sleep spindle density (number of spindles per 30 s) from all detected spindles.

### Slow Wave Detection

We used YASA’s slow wave detection algorithm that is based on Carrier [40] and Massimini and colleagues [41]. YASA first filters the signal from 0.3 to 3.5 Hz. Next it finds all negative peaks with an amplitude between -40 to -300 µV and all positive peaks with an amplitude between 10 to 200 µV. If the amplitude from negative peak to positive peak is between 75 to 500 µV, several duration criteria apply, including a negative-deflection-phase duration between 0.3 to 1.5 s and a positive-deflection-phase duration between 0.1 to 1 s. We continued our analyses with slow waves detected on channel Fpz, which was used for auditory stimulus targeting, computing duration and peak-to-peak amplitude for each detected slow wave. We also estimated slow wave density (number of slow waves per 30 s) from all detected slow waves.

### Statistical Analysis

Possible differences between the stimulation and sham condition for PSDs, ERPs, TFRs, as well as stimulus-locked average number of slow waves and spindles, and slow wave – spindle coupling, were assessed through mass-univariate statistical comparisons, using threshold-free cluster enhancement (TFCE) permutation testing methods [42], as implemented in the open source Python package Eelbrain [43]. For general information on this method see the Supplementary Information. Within-subjects measurements were compared with a TFCE using F-values resulting from repeated measures ANOVA (TFCE_ANOVA_). We then determined p-values by running 10,000 permutations and finding the fraction of TFCE values at least as extreme as the original TFCE value. Significant clusters were detected by thresholding p-values with a significance level of 𝛼 = 0.05. Post-hoc pairwise comparisons with TFCE were conducted based on paired t-tests (TFCE_t_); significant clusters were found with a Holm-Bonferroni correction to account for multiple pairwise comparisons. Finally, only time-frequency-channel triplets that were significant in the TFCE_ANOVA_ and the TFCE_t_ were allowed to form clusters.

Given our experimental design, the analysis of short- and long-term effects of auditory stimulation on sleep spindle and slow wave characteristics (spindles: amplitude, duration, frequency and density; slow waves: amplitude, duration and density) required methods that can handle repeated-measures, non-normal data, as well as unbalanced observations arising from the inter-subject variability in sleep duration and the randomized presentation of conditions. Linear mixed effects models (LMMs) or generalized linear mixed effects models (GLMMs) can deal with these circumstances. Thus, after checking our assumptions about the observations’ distributions, they were used where appropriate.

We chose the fixed effects structure of our model *a priori* based on our hypotheses. We included a fixed effect “Condition”, with levels corresponding to the stimulation conditions (STIM, SHAM), and another fixed effect “Term”, describing the temporal relationship to stimulus onset, with levels referencing to either short-term (SHORT) or long-term (LONG). Throughout the remainder of this article short-term refers to the stimulation-epoch from 0 to 3 s after stimulus onset and long-term corresponds to the rest of the CB (i.e. the periods between subsequent stimulation-epochs or between such epochs and the end of a CB; long-term periods had a median duration of 5.3 s; for further characteristics of long-term periods see Supplementary Information and Supplementary Figure 1).

Initially, we specified all models with the maximum random effects structure justified by the experimental design [44]. Next, we incrementally simplified models not converging with the maximum random effects structure or if overfitting was present (see Supplementary Tables 1 and 2 for the final structure of each model). All models were fitted with the afex::mixed R package (Henrik Singmann, Ben Bolker, Jake Westfall, Frederik Aust and Mattan S. Ben-Shachar, 2019), using (restricted) maximum likelihood estimation. Random effects were integrated out using the Laplace approximation. After fitting the models, we did several visual and statistical model diagnostics such as residual plots, quantile-quantile plots and Kolmogorov-Smirnov tests using the simulation-based R package DHARMa (Florian Hartig, 2019). We conducted hypothesis tests on the fixed effects with either the Kenward-Roger approximation using F-tests for LMMs, or by comparing the original likelihood-ratio to a reference distribution generated by parametric bootstrapping (PB) using 10,000 simulations for GLMMs. If fixed effects or their interactions were found to be significant, we ran post-hoc tests on their estimated marginal means (EMMs) with multiple comparison adjustments based on multivariate-t-distributions using the R package emmeans (Russell Lenth, 2019). As main effects of the Term factor were not of primary interest to our objectives, they were not reported. Differences between factor levels are reported as EMMs contrast ± standard error.

## Results

### Sleep architecture

On average, participants slept for 90.09 minutes (SD = 30.98) during the experiment, of which 41.9% (SD = 19.5) was NREM 3 sleep (Table 1).

**Table 1.** Polysomnographic Results.

|  | ***Total sleep***<br>***time (min)*** | ***NREM 1***<br>***(% TST)*** | ***NREM 2***<br>***(% TST)*** | ***NREM 3***<br>***(% TST)*** | ***REM***<br>***(% TST)*** |
| --- | --- | --- | --- | --- | --- |
| ***Mean*** | 90.09 | 15.8 | 30.9 | 41.9 | 11.4 |
| ***Std. dev.*** | 31.0 | 7.2 | 12.8 | 19.5 | 10.9 |

*Note.* This table shows the polysomnographic sleep staging results of the afternoon nap. TST = Total Sleep Time; min = minutes; %TST = percentage of the total sleep time, NREM = non-REM, std. dev = standard deviation,

### Phase-targeting accuracy

As a manipulation check, we evaluated the phase-targeting performance of our closed-loop phase prediction method. The circular mean and standard deviation of the slow wave phase at the onset of sound stimuli, at Fpz, across all participants and conditions, were 5.92 ± 52.83° (Figure 1B). Thus, our method was able to target the intended slow oscillation phase (0°) consistently, with high precision.

### Amplitude-modulated white noise and white noise stimulation

As a first analysis step, we investigated whether AMWN and WN elicited distinct neural responses. As we expected sleep spindles might acutely entrain to AMWN, we studied ERPs and TFRs in the 3 seconds following stimulus onset. We also investigated discrete sleep spindle induction due to AMWN and WN by analyzing sleep spindle density, considering both short-term and long-term responses.

Our findings indicated no differential effects of the two distinct auditory stimulation conditions (see Supplementary information). Hence, the AMWN and WN data were pooled for further analyses. From here on, these combined data will be referred to as the stimulation condition: STIM.

### Power spectral density

To assess overall effects of auditory stimulation on sleep’s spectral composition, we analyzed spectral differences between the STIM and SHAM condition, in CB blocks pooled per condition, across a broad frequency range, from 0.1 to 40 Hz (Figure 2A). Comparing PSDs in STIM versus SHAM returned three significant clusters, one from 0.2 to 1.6 Hz (cluster p = 0.0023), another from 2.1 to 2.4 Hz (cluster p = 0.0364) and the third from 11.6 to 14 Hz (cluster p = 0.002). The first two clusters resided in the slow wave and delta range, respectively, and reflect increased power in the auditory stimulation condition. We found that frontal channels, expressing a commonly observed slow wave topography, were significantly involved in driving power changes (Figure 2B, top). The third cluster partially covered the sleep spindle range and reflects reduced power with auditory stimulation. This cluster showed significant power changes in most channels with pronounced effects in centroparietal and occipital regions consistent with fast sleep spindle topographies (Figure 2B, bottom). These results suggest that auditory stimulation boosts slow wave and delta activity, while suppressing spindle activity throughout CBs.

**Figure 2.**
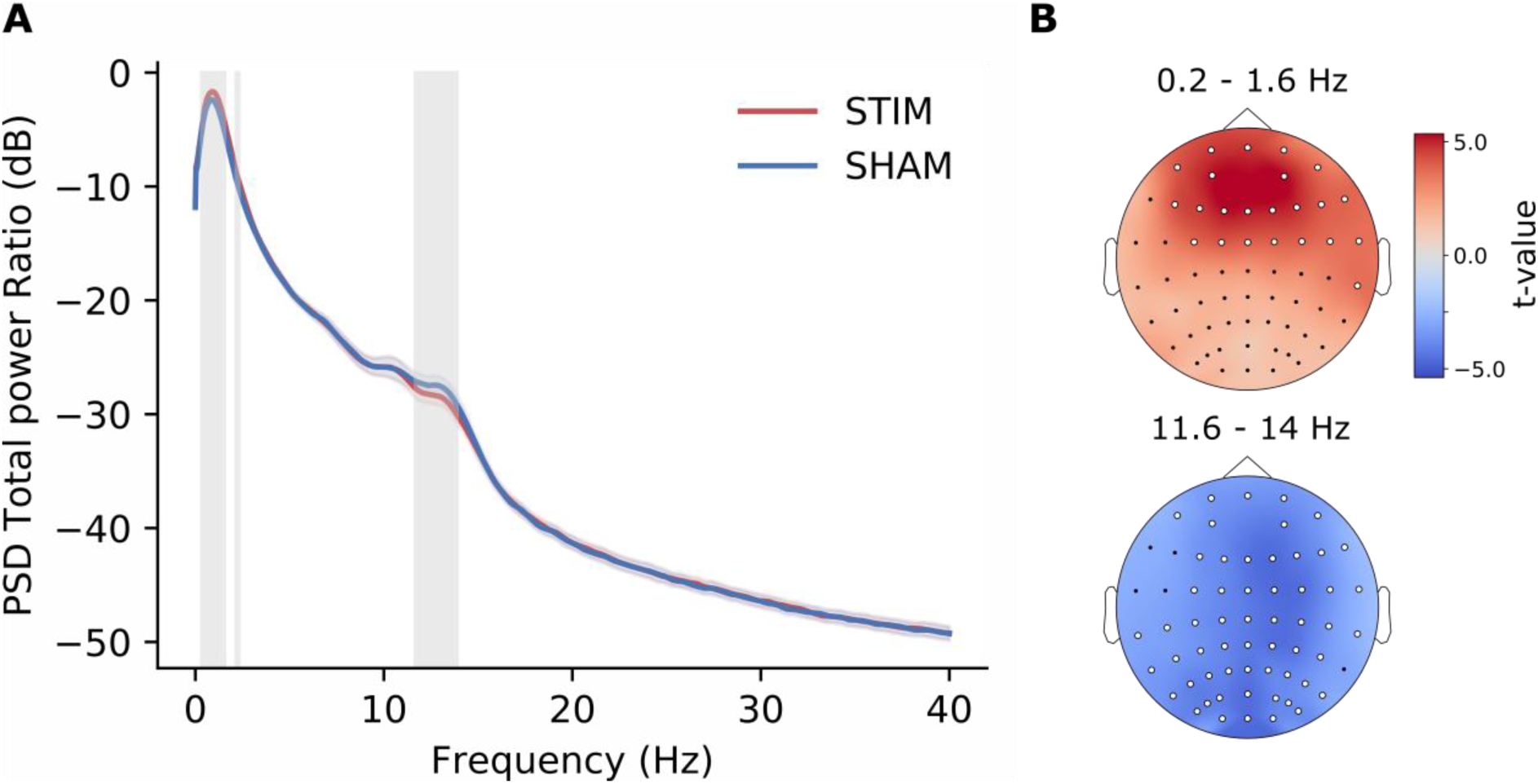
Power Spectral Density. (A) Plotted are the power spectral density estimates of STIM and SHAM (total power normalized) with their respective SEMs for channel Fpz. The gray shaded areas indicate significant TFCEt clusters based on all channels. (B) Shown are the topographic differences (t-value) between STIM and SHAM for the slow wave cluster (0.2 to 1.6 Hz) and the spindle cluster (11.6 to 14 Hz). Significant channels are indicated with white markers. The second cluster, possibly reflecting delta or slow wave activity, only involved five frontal channels that were also part of the slow wave cluster and is not shown.

### Acute dynamics of sleep spindles and slow waves in response to auditory stimulation

Next, we studied the acute effects of auditory stimulation, aiming in particular to understand how responses of discrete slow waves and sleep spindles may relate to effects at the level of ERPs and TFRs. To start, we compared the ERP-responses for STIM and SHAM. The STIM ERP shows a sequence of more pronounced slow positive and negative deflections compared to the SHAM ERP, leading to three significant clusters (Figure 3A). Referenced to stimulus onset (0 s), the first cluster (cluster p = 0.0001) occurs from 0.3203 to 0.7676 s, the second (cluster p = 0.0001) from 0.8457 to 1.2422 s, and the third (cluster p = 0.0024) from 1.5078 to 1.8711 s. These results could reflect an increase of amplitude or occurrence of slow oscillations, and/or a more stereotypical shape of stimulation-evoked slow wave activity leading to enhanced phase-alignment. Of note, the ERP period between -1 and 0 shows a slow oscillation-like deflection, which crosses time zero at the expected 0° phase, for both the STIM and SHAM conditions. This corroborates the algorithm’s success at detecting SOs and targeting 0°.

**Figure 3.**
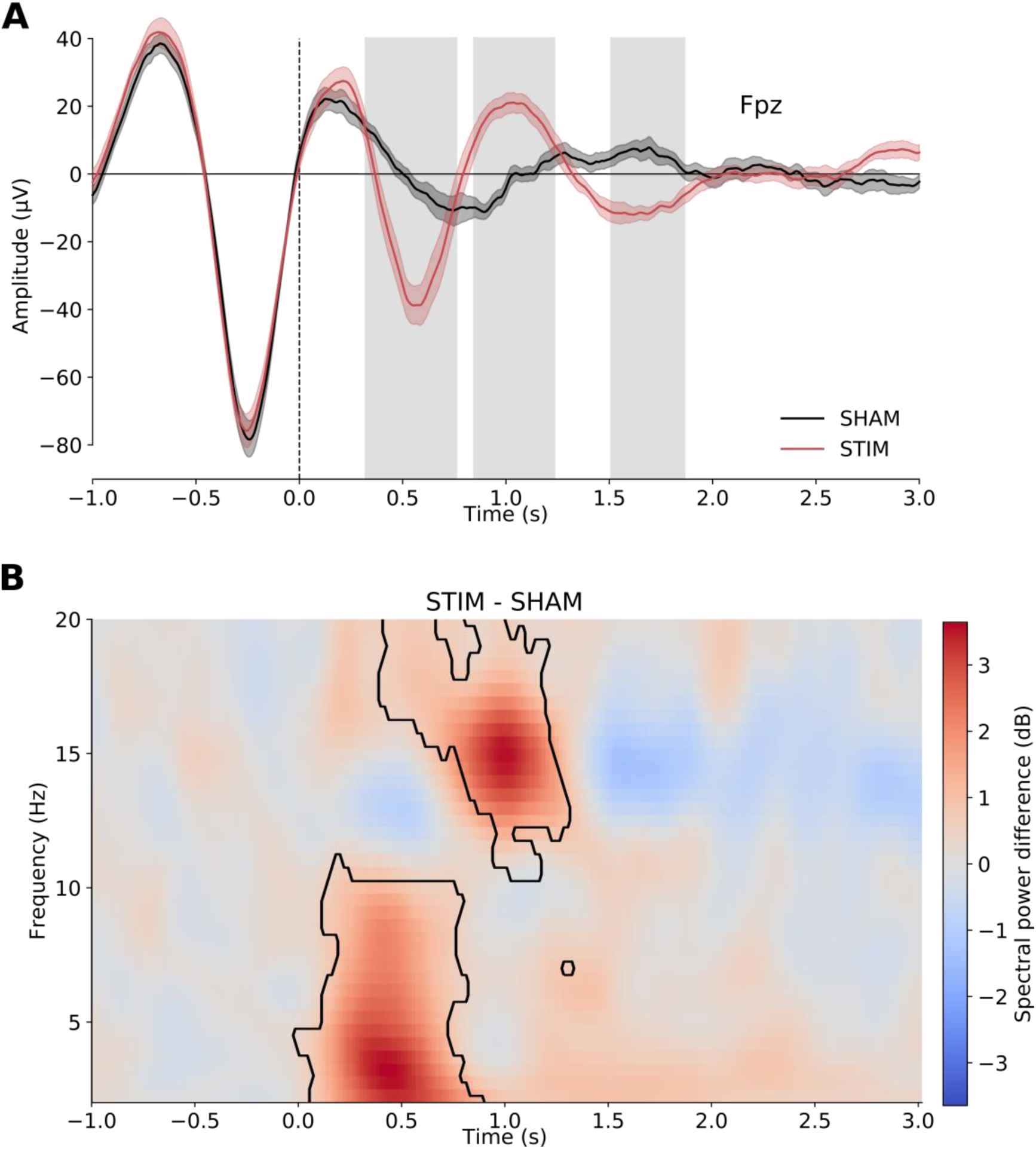
Event-related potentials and time-frequency representation. (A) Plotted are the grand averages of the STIM and SHAM conditions with their respective SEMs for channel Fpz. The gray shaded areas indicate significant TFCEt clusters based on all channels. (B) The spectral power differences in dB between STIM and SHAM are color-coded. Red color indicates higher power in STIM, while blue indicates higher power in SHAM. Clusters where the spectral power differences were significantly different (obtained by TFCEt) are marked by a black outline.

A subsequent TFR analysis, comparing STIM to SHAM, revealed two significant clusters (Figure 3B). One early cluster, from 0 to 0.9023 s referenced to stimulus onset (0 s), is in the slow wave/delta/theta range (2 to 11 Hz, cluster p = 0.0001); the other later cluster, from 0.3945 to 1.3125 s, is in the sigma/beta range (10.5 to 20 Hz, cluster p = 0.0021). The first cluster (2 -11 Hz) suggests that auditory stimulation, in agreement with previous findings [5–9,17], may induce K-complexes or slow waves, and/or boosts their peak-to-peak amplitudes, and/or redistribute their occurrence along the considered time window. The second cluster suggests the same for spindle/beta activity. Broadly speaking, the ERP and TFR findings parallel previous observations [6–9].

We next turned to analyses of discrete slow oscillation and sleep spindles to understand the effects of stimulation on these oscillatory events in more detail. We asked whether the enhanced slow deflections in the ERPs and increased low-frequency power cluster in the TFRs might reflect increased occurrence of discrete slow oscillations. Similarly, we assessed if fast spindles co-occurring with boosted and/or induced slow wave positive deflections could drive the frequently observed power increase in the TFR sigma cluster following auditory stimulation. To address these questions, we continued by establishing the relationship of discrete sleep spindles and slow waves to ERPs and TFRs.

We first characterized the temporal dynamics of discrete slow wave positive deflections by analyzing their distribution across stimulation epochs (-1 to 3 s referenced to stimulus onset). To this end, epochs were split into time-bins (200 ms); the number of slow wave positive deflections peaks in Fpz, normalized by the total number of epochs within each stimulation-condition, was determined for each bin. We found a significant increase of the average number of slow wave positive deflections peaks in STIM compared to SHAM in the bin from 0.8 to 1.0 s after stimulus onset (SHAM vs. STIM: t(13) = -4.1911, p = 0.0168; Figure 4A). Notably, increased occurrence of slow wave positive deflections aligns with the second ERP-cluster and the TFR-sigma-cluster (Figure 3A and B).

**Figure 4.**
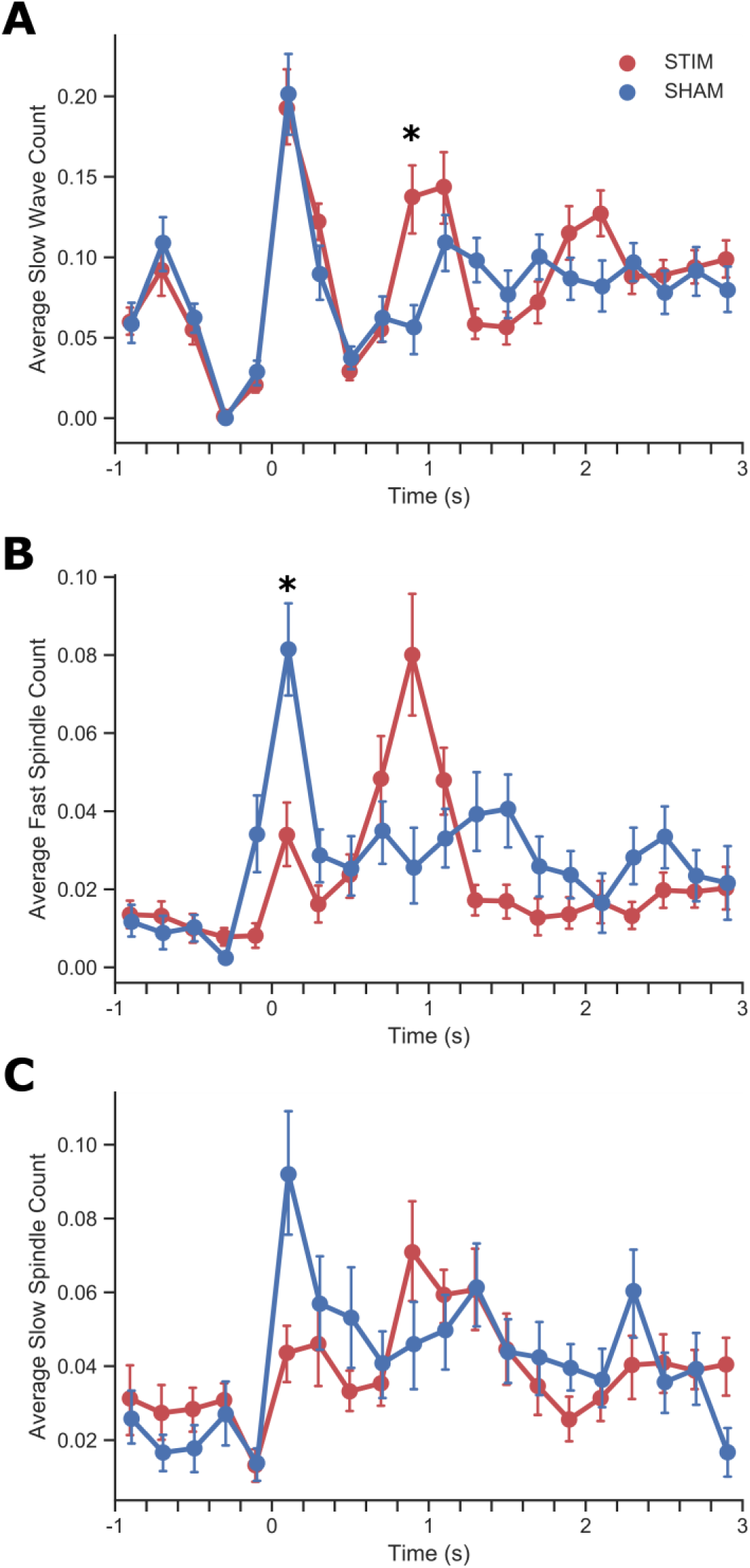
Acute dynamics of discrete spindles and slow waves in response to auditory stimulation. Distributions were created by computing the average number of events per bin (length of 200 ms) per stimulus across epochs (time is referenced to stimulus onset at 0 s). The circular markers indicate the mean across all participants (n = 14) and the bars represent the SEM. Significant differences between STIM and SHAM bins are indicated by asterisks. (A) Distribution of average number of slow waves (referenced to slow wave positive deflection peaks). (B) Distribution of average number of fast spindles (referenced to spindle peak). (C) Distribution of average number of slow spindles (referenced to spindle peak)

Following the same procedure as for slow waves, we next investigated the distribution of discrete fast and slow sleep spindles across epochs (sleep spindles were detected across all channels). Contrary to our expectations, the average number of fast spindles only differed significantly between stimulation conditions in the first bin, with fewer fast spindles following auditory stimulation (t(13) = 4.8349, p = 0.0005; Figure 4B). However, auditory stimulation led to a non-significant peak in the number of fast spindles from 0.8 to 1 s (SHAM vs. STIM: t(13) = -2.7152, p = 0.1913), aligned with the increase in slow wave positive deflections (Figure 4A), the second ERP-cluster (Figure 3A), and the TFR-sigma-cluster (Figure 3B). In contrast to fast spindles, slow spindle occurrences were not significantly affected by the auditory stimulation (Figure 4C). These findings suggest a 0°-targeted auditory stimulus leads to enhanced occurrence of slow oscillations and an immediate, very brief disruption of fast spindles or their generation processes.

### Slow wave – spindle coupling

Because previous studies have suggested a relationship between slow wave – spindle coupling and memory retention, we studied the effects of auditory stimulation on the coupling of slow waves with discrete fast and slow spindle occurrences. For this analysis, we counted spindles (across all channels) occurring in time-bins (200 ms) from -1 to 1 s referenced to the peak of slow waves’ negative deflections (SWND; in Fpz) and normalized the results by the number of slow wave – spindle couplings within the respective stimulation-condition (Figure 5). We did not detect significant differences, either for fast (Figure 5A) or slow spindles (Figure 5B), across any of the slow wave phases. Therefore, auditory closed-loop stimulation appears to have no significant effect on slow wave – spindle coupling.

**Figure 5.**
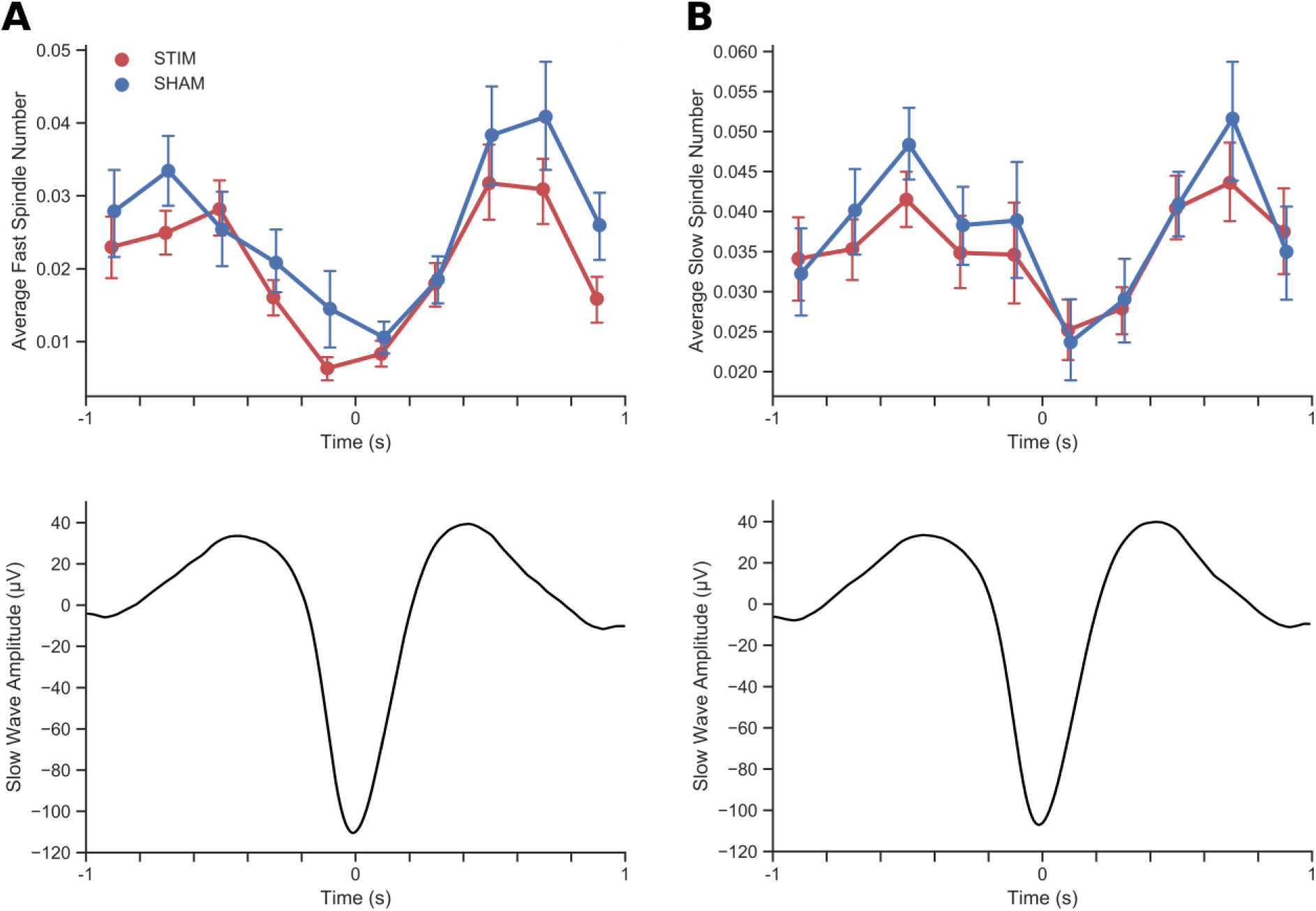
Slow wave – spindle coupling. The distributions were created by computing the average number of spindle events per bin (length of 200 ms) per slow wave (time is referenced to slow wave negative peak at 0 s). The ciruclar markers indicate the mean across all subjects (n = 14) and the bars represent the SEM. The top graphs in (A) show the average fast spindle number per bin per slow wave for STIM (red) and SHAM (blue). No significant differences between the two conditions could be detected. The bottom figure shows the average of fast spindle hosting slow waves across all subjects and conditions in channel Fpz. The top figure in (B) shows the average slow spindle number per bin per slow wave for STIM (red) and SHAM (blue). No significant differences between the two conditions could be detected. The bottom figure shows the average of slow spindle hosting slow waves across all subjects and conditions in channel Fpz.

### Short-term and long-term effects of M-CLNS on slow waves

As one of our main objectives, we then investigated the short-term and long-term responses of discrete slow waves to auditory stimulation (Figure 6). We considered density, amplitude and duration of slow waves.

**Figure 6.**
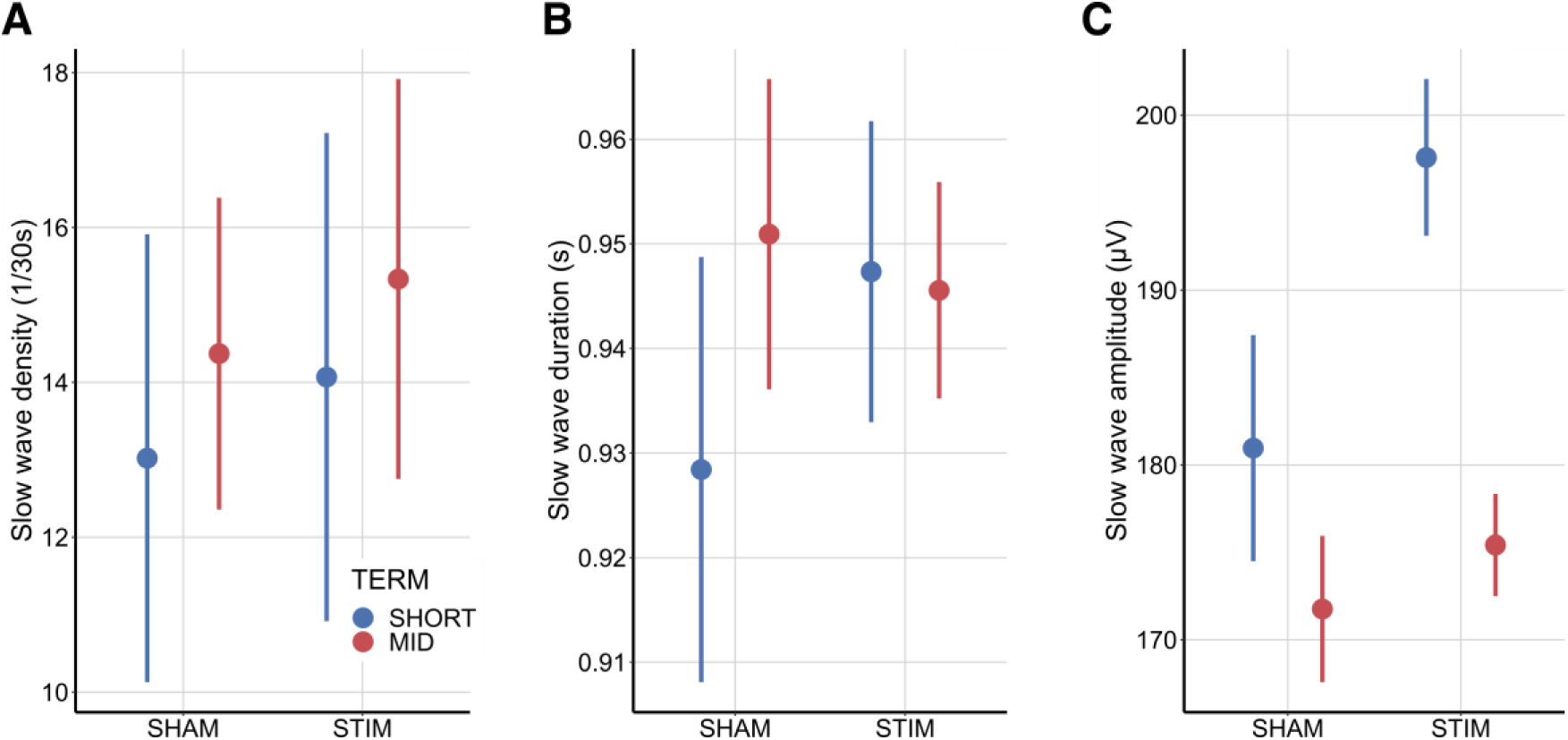
Individual slow wave characteristics. This figure shows the descriptive data means (circular markers) and their respective 95% confidence intervals alongside the EMMs (diamond markers) split by Condition and Term. In some instances, the confidence intervals are covered by the markers. In the background we show the distribution of individual datapoints (not transformed). (A) depicts the slow wave density. (B) shows the slow wave duration. (C) shows the slow wave amplitude.

Our analysis revealed a trend for increased slow wave density by auditory stimulation (STIM vs SHAM: 1.0053 ± 0.5147 Slow waves/30s, CI_95%_ = [0.1065, 2.117]; F(1,13) = 3.8157, p = 0.0726). The Condition x TIME interaction (F(1,13) = 0.0077, p = 0.9315) had no significant effect (Figure 6A). In addition, all predictors and their interaction significantly affected slow wave amplitude (Condition: F(1,13.104) = 37.086, p < 0.0001; Condition x Term: F(1,7036.11) = 17.804, p < 0.0001) (Figure 6C). In the short-term slow wave amplitude was 22.34 ± 3.39 µV higher during STIM compared to SHAM (CI_95%_ = [13.63, 31.0]; z = 6.589, p < 0.0001), while the amplitude was elevated by 6.36 ± 2.52 µV (CI_95%_ = [0.12, 12.8]; z = 2.521, p = 0.0427) in the long-term. Finally, auditory stimulation had no significant effect on slow wave duration (Condition: F(1,7064.46) = 0.2441, p = 0.6213; Condition x Term: F(1,4997.05) = 1.9021, p = 0.1679) (Figure 6B). These findings show that auditory stimulation clearly enhances slow waves both on the short- and long-term, mostly in terms of amplitude.

### Short-term and long-term effects of M-CLNS on fast spindles

Similar to slow waves, we investigated the short-term and long-term responses of fast spindle density, amplitude, duration and frequency to auditory stimulation.

We found that fast spindle density was 1.0365 ± 0.2996 spindles/30s lower during auditory stimulation compared to sham (CI_95%_ = [0.3892, 1.6839]; F(1,13) = 11.9678, p = 0.0042). The Condition x Term interaction (F(1,13) = 0.3924, p = 0.5419) had no significant effect (Figure 7A). Fast spindle duration was also lower during auditory stimulation compared to sham, by 25.1 ± 8.2 ms (CI_95%_ = [9.06, 41.2]; 𝜒^2^(1) = 10.3958, PB p = 0.0029). The Condition x Term interaction (𝜒^2^(1) = 0.2669, PB p = 0.6120), again, had no significant effect (Figure 7B). These drops in fast spindle density and duration due to the condition main factor might be related to the earlier shown CB-wide increase of slow wave activity due to auditory stimulation.

**Figure 7.**
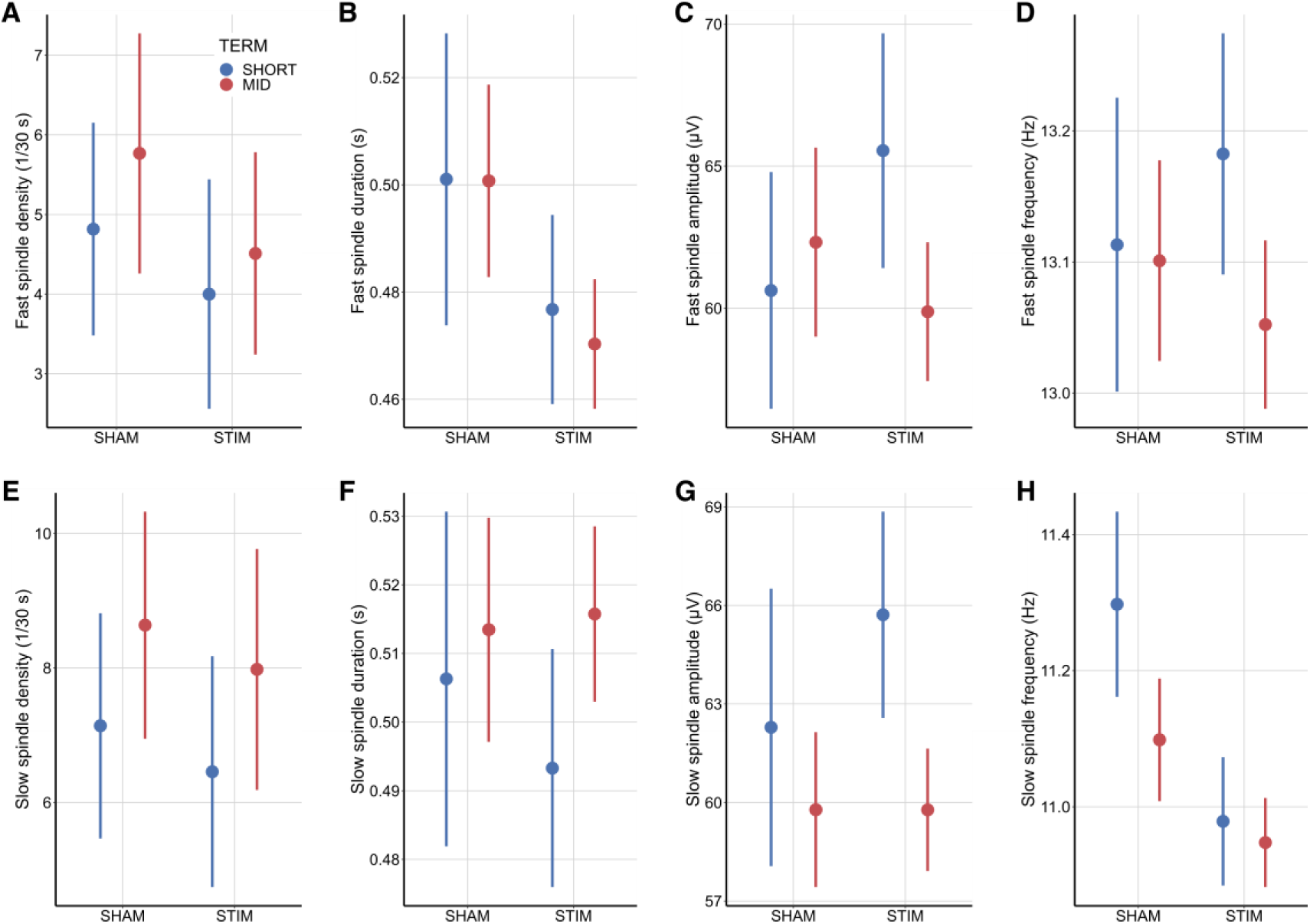
Individual spindle characteristics. The figures show the descriptive data means (circular markers) and their respective 95% confidence intervals alongside the EMMs (diamond markers) split by Condition and Term. In some instances, the confidence intervals are covered by the markers. In the background we show the distribution of individual datapoints (not transformed). (A) depicts the fast spindle density. (B) shows the fast spindle duration. (C) shows the fast spindle amplitude. (D) shows the fast spindle frequency. (E) depicts the slow spindle density. (F) shows the slow spindle duration. (G) shows the slow spindle amplitude. (H) shows the slow spindle frequency.

Fast spindle amplitude, showed a significant influence of the Condition x Term interaction (F(1,2178.9) = 4.0888, p = 0.0433), while Condition (F(1,2180.9) = 1.6266, p = 0.2023) alone had no significant effect (Figure 7C). Post-hoc tests, showed that in the long-term fast spindle amplitude was 3.621 ± 1.26 µV lower during auditory stimulation compared to sham (CI_95%_ = [-0.0463, 7.288]; t(14) = 2.871, p = 0.0414), while no significant short-term effect was detected (SHORT-SHAM vs SHORT-STIM: -0.824 ± 1.85 µV, CI_95%_ = [-6.1309, 4.482]; t(16) = -0.444, p = 0.9609). Furthermore, we found no significant differences between short-term and long-term effects within stimulation conditions (SHAM-MID vs SHAM-SHORT: 1.241 ± 1.76 µV, CI_95%_ = [-3.8300, 6.311]; t(14.8) = 0.706, p = 0.8668; STIM-MID vs STIM-SHORT: -3.205 ± 1.38 µV, CI_95%_ = [-7.2183, 0.809]; t(14) = -2.321, p = 0.1117).

Fast spindle frequency was not significantly affected by Condition (F(1,2059,337) = 0.0265, p = 0.8706) nor Condition x Epoch (F(1, 1820.241) = 0.7668, p = 0.3813) (Figure 7D).

### Short-term and long-term effects of M-CLNS on slow spindles

With the same procedure as outlined for fast spindles, we studied the short-term and long-term responses of slow spindle density, duration, amplitude, and frequency to auditory stimulation.

We found no significant effect of auditory stimulation on slow spindle density (Condition: F(1,26) = 2.7783, p = 0.1076; Condition x Term: F(1,26) = 0.0008, p = 0.9779; Figure 7E), duration (Condition: 𝜒^2^(1) = 0.5892, PB p = 0.4441; Condition x Term: 𝜒^2^(1) = 0.9309, p = 0.3382; Figure 7F), or amplitude (Condition: F(1,13.887) = 0.0066, p = 0.9362; Condition x Term: F(1,14.511) = 0.5462, p = 0.4717; Figure 7G).

Slow spindle frequency was the only characteristic affected and was 0.156 ± 0.0306 Hz lower during auditory stimulation compared to sham (CI_95_ = [0.0963, 0.216]; F(1, 15.438) = 25.5346, p = 0.0001) . The Condition x Term interaction (F(1,3376.567) = 0.9099, p = 0.3402) was not significant (Figure 7H).

## Discussion

In this study we assessed a novel modelling-based closed-loop stimulation (M-CLNS) procedure with accurate and precise slow wave phase-targeting, and single rather than rhythmically spaced sound pulses. As a secondary objective, we investigated whether pulses of white noise with amplitude modulation in the spindle frequency range could be especially effective in modulating sleep spindles. We hypothesized that the M-CLNS method might have a more enduring effect on oscillatory sleep dynamics compared to methods used thus far. We investigated short-term effects of individual stimuli, as well as overall effects across stimulated sleep. We include a detailed analysis of discrete fast and slow spindles, which have not been extensively evaluated thus far in the field of closed-loop acoustic sleep stimulation.

With respect to our objective of studying short- and longer-term responses of slow waves and sleep spindles to the stimulation protocol, our main finding is that slow wave measures were consistently increased, while fast spindle measures were consistently decreased. These changes were globally reflected in PSD differences between auditory and sham stimulation in the slow wave, delta, and sigma bands. The slow wave and delta effects were localized to a large frontal cluster, while the sigma band effects covered most of the scalp. More detailed analyses on discrete slow wave and spindle events confirmed and extended these findings.

Starting with slow waves, auditory stimulation led to a considerable amplitude increase in the short-term and a smaller increase in the long-term, accompanied by a global trend towards increased density of slow waves (∼2 slow waves more per minute). Fast spindles, on the other hand, showed a global reduction in density (∼2 spindles less per minute) and duration. Additionally, fast spindle amplitude decreased in the long-term. Slow spindles were less consistently affected by auditory stimulation, although there was some tendency toward reduced density and a significant reduction in frequency by auditory stimulation.

Analysis of the first 3 seconds following stimulus onset showed that the acute dynamics of discrete slow wave and fast spindle events were significantly altered by auditory stimulation: slow wave positive deflections occurred more frequent around 1 second after auditory stimulation compared to sham; this increase was aligned with a significant positive deflection in the ERPs and a significant sigma/beta cluster in the TFRs. Fast sleep spindle occurrences were reduced in the 200 ms after stimulus onset, suggesting sounds disrupted ongoing spindle dynamics and/or the momentary generation of spindles. Notably, our findings showed no differential effects between standard white noise and amplitude-modulated white noise, which was specifically intended to enhance spindles. These results suggest that amplitude modulation in the spindle frequency has no added effects on spindle modulation.

To our knowledge, this is the first study to demonstrate that acoustic stimulation leads to enduring sleep deepening by presenting single acoustic stimuli precisely phase-locked to slow oscillations. This effect is evident from several measures taken across stimulated sleep, which show increased power in the SO and delta frequency ranges, increased SO amplitude and a trend-level increase in SO density. This finding adds to the literature, in which mostly rhythmic stimulation was used [9,19,20,45], leading either to no overall sleep deepening or an enhancement of power around the stimulation frequency (e.g. 1 Hz), potentially reflecting an artifact of rhythmic stimulation. In support of this perspective, studies have shown that the timing of stimuli relative to the ongoing slow oscillation influences the response [6,18] and that untimed stimuli can disturb traveling SO waves [22]. Moreover, responses to rhythmic stimulations were shown to die out quickly, possibly because untimed stimuli in the sequence interfere with the natural oscillation dynamic [5,8,21]. In contrast, our findings show that single, phase-locked acoustic stimuli precisely targeted to the rising phase of slow oscillations are able to induce enduring enhancements of SO dynamics.

While acoustic stimulation enhanced slow oscillations, it also globally reduced sleep spindles. This may be explained by the inverse relationship between sleep depth and spindles, where the occurrence of especially fast sleep spindles decreases with increasing sleep depth [46].The global fast spindle reduction may thus be viewed as supporting evidence for the sleep deepening effects of acoustic stimulation.

While long-term sleep spindles were reduced, short-term sigma power was increased in a short time window after acoustic stimulation, as shown by time frequency responses. The sigma increase coincides temporally with enhanced or induced SO positive deflections around 1 second after stimulation. In this same time window, we observed a non-significant but pronounced occurrence of discrete fast spindles, suggesting the sigma increases might reflect fast spindles. These findings are consistent with previous studies reporting similar sigma power increases time-locked to acoustic stimulation [5–9,16,17,47]. Factors potentially contributing to the stimulus-locked spindle increase are, first, a more temporally consistent response of neural activity to the stimulus, compared to spontaneous activity (in the SHAM condition), leading to higher temporal alignment (across trials) of both slow oscillation dynamics and the spindle cluster [6,47]. Secondly and more speculatively, the spindle suppression at stimulus onset might lead to a rebound of spindle activity at the next permissive moment, i.e. the next SO-depolarizing upwave [48,49].

Several limitations should be considered when interpreting these findings. The sleep EEG in this study was obtained from of a daytime nap. Caution is warranted when generalizing these findings to nocturnal sleep. The body’s physiological state may vary between day- and nighttime sleep due to differences in both central and peripheral circadian rhythms [50]. Future studies could adopt an overnight design to assess its impact on the physiological, restorative aspects of sleep.

Another limitation is that this sample primarily consisted of healthy, young subjects, as is the case in the majority of acoustic sleep stimulation studies [6,23,51,52]. Other age groups for which slow oscillation enhancement in response to acoustic stimulation has been observed are children (8-12 years) [53] , middle-aged men (35-48 years) [54] and older adults (60-84 years) [7]. Although the specific phase-targeting algorithms for slow oscillations vary across these studies and should therefore not be viewed as a single method [7,23,53,54], the cumulative findings suggest that SO enhancement has strong potential for widespread application throughout the lifespan. Given that slow wave sleep decreases with age [55,56], interventions targeting sleep deepening may be especially beneficial for older adults.

The amplification of slow oscillations may offer therapeutic potential for patient populations with insufficient deep sleep, including those with insomnia [57] and other sleep disorders [58–60]Sleep enhancement may also be valuable in patients with mental disorders, including post-traumatic stress disorder [61,62], depression [63], schizophrenia [64] , and anxiety disorders [62] considering that sleep disturbance predisposes for such disorders [65–67], and exacerbates clinical symptom severity [68–70]. More generally, restoring deep sleep is important given to its vital role in a wide range of restorative processes that together protect brain function [71], cardiovascular health [72], immune responses [15,73], and reduce the risk of metabolic disorders, like type 2 diabetes, through glucose homeostasis [74].

In addition to the potential for deepening sleep with acoustic stimulation of slow oscillations, memory enhancement effects have also been widely reported, both in healthy individuals [14,17,75–77], and in patients with mild cognitive decline [78]. Also here, the timing of acoustic memory cues relative to spontaneous slow oscillation [79] and spindle dynamics appears to matter [49]. Notably, fast spindle activity has been widely implicated in memory reprocessing during sleep [48,80]. Hence, the synchronized increase in fast spindle occurrence shortly after stimulus onset, observed in our study might well play a role in reprocessing memories [48,80]. To optimize the memory enhancement effects of closed-loop acoustic stimulation, the stimuli may be presented in response to spindle detection [81,82] rather than detecting slow oscillation.

In summary, we found that slow oscillations were enhanced both in the short and longer term in response to single, phase-locked acoustic stimuli, while spindles were reduced. Future research may explore the extension of slow oscillation enhancement in a multi-night design. Although this study focused on a young healthy cohort, further research on M-CLNS could elucidate its potential effectiveness throughout the lifespan and in patient populations suffering from sleep disorders.

## Supporting information

Supplemental Results

## Acknowledgements

We thank the European Space Agency (ESA) and Leopold Summerer for their support and collaboration on this study. Contract number: 4000125903/18/NL/GLC/as.

