## Supplemental Results for "Precisely phase-locked acoustic stimuli globally enhance slow oscillations, but depress fast spindles"

### Supplementary Information

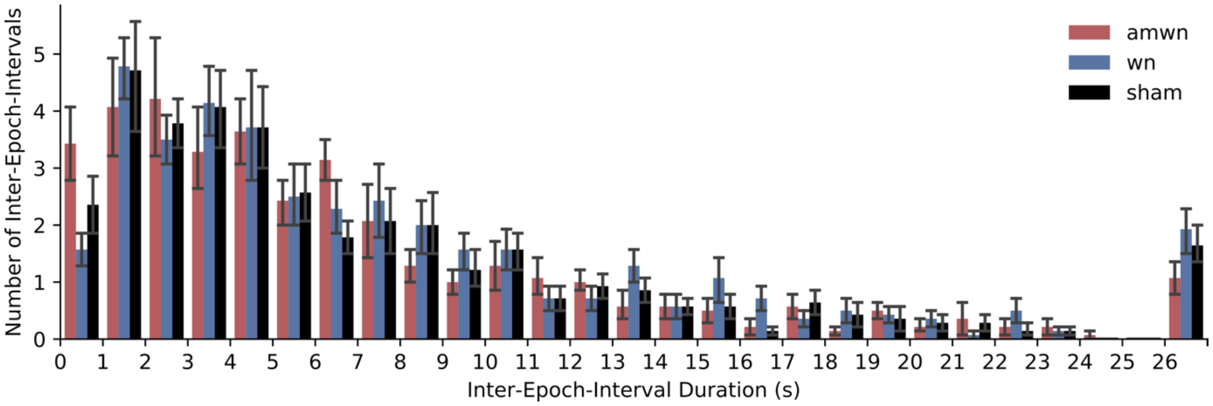


*Supplementary Figure 1 – Distribution of Inter-Epoch-Intervals (mid-term periods). An Inter-Epoch-Interval (IEI) is defined as the period between subsequent stimulation-epochs (0 to 3 s after stimulus onset) or the end of a condition block. The bars indicate the average number of IEIs in a specific bin (width of 1 s) across all participants and blocks, and the error bars indicate the SEM. $TFCE_{ANOVA}$ did not detect any significant differences in the distributions of IEIs between conditions (AMWN, WN, SHAM). The distributions are described by a median of 4.95 s ($Q_1$ = 2.32 s, $Q_3$ = 9.20 s) for AMWN; a median of 5.85 s ($Q_1$ = 3.00 s, $Q_3$ = 10.61 s) for WN, and a median of 5.05 s ($Q_1$ = 2.53 s, $Q_3$ = 9.94 s) for SHAM. The increase of IEIs in the 26-27 s-bin represents all the CBs that only had one stimulation at the beginning, making the duration of 27 s of the subsequent IEIs non-random.*

*Threshold-free cluster enhancement*

TFCE controls for multiple comparisons that arise from mass-univariate testing by taking individual voxels' correlations into account. As opposed to standard cluster-based permutation tests, TFCE avoids researcher bias introduced by threshold selection and, furthermore, allows interpretation of individual voxels.

First, a statistic (e.g. F- or t-value) is calculated for each voxel between conditions. Based on the resulting statistic landscape the TFCE value of an individual voxel is computed by integrating from zero to the voxel's statistic value, i.e. summing all potential thresholds, while multiplying the value by the number of adjacent voxels that reach at least the same threshold. Avoiding the continuous integral notation and instead using the more practical sum notation the TFCE value for e.g. time-frequency-channel triplets (for simplicity denoted with i) can be described as

$$TFCE(i) \ = \sum_{0}^{h(i)} e^{E}\ (i)\ h^{H}(i)$$

Here $h(i)$ denotes a value that is increased from zero to the statistic value of the triplet, $e(i)$ refers to the number of neighbouring triplets that reach at least the potential threshold $h(i)$, and E and H are parameters that are set to 0.5 and 2.5 on theoretical grounds (for details see Smith & Nichols, 2009). Intuitively the TFCE enhances statistic values that have strong intensity and/or are supported by many neighboring voxels while preserving local extrema of the statistic landscape. This latter property allows for interpretation of extrema within clusters. Last, a null distribution of TFCE values is created for each voxel using permutation tests. The p-value of a voxel corresponds to the fraction of TFCE values in the null distribution that are at least as extreme as the original TFCE value.

*Distribution of mid-term periods.* Characteristics of slow waves and spindles were assessed both short-term within stimulation-epochs (0 to 3 s after stimulus onset) and mid-term within inter-epoch-intervals (IEIs), which were defined as periods between subsequent stimulation-epochs or the end of a condition block ("Term"-predictor with the level "SHORT" corresponding to within stimulation-epochs and "MID" corresponding to within IEIs). The IEI-distribution across all participants and stimulation-conditions had a median of 5.30 s (Q1 = 2.59 s, Q3 = 10.07 s) and a minimum and maximum of 0.05 and 27.00 s, respectively. To quantify if IEIs are structured by the stimulation-conditions we conducted a $TFCE_{ANOVA}$ on the IEI-distributions (Supplementary Figure 1), which indicated no significant effects.

*Amplitude-modulated white noise and white noise stimulation have no differential effects*.
First, we analyzed whether the condition (AMWN, WN, SHAM) influenced the number of occurring stimulations. Since the number of stimulations was not normally distributed, we conducted a Friedman test to compare the effect of the stimulation-condition. The test did not show any significant differences between the number of stimulations in each condition ($\chi^2(2) = 1$, p = 0.6065).

Comparing ERPs between AMWN, WN, and SHAM stimulation using $TFCE_{ANOVA}$, revealed three significant clusters from 0.334 to 0.756 s (cluster p < 0.0001), 0.840 to 1.201 s (cluster p = 0.0002) and 1.52 to 1.830 s (cluster p = 0.0016). Post-hoc $TFCE_t$ comparisons showed that those differences were driven by both AMWN and WN differing from the SHAM condition (Supplementary Figure 2A), while the two auditory conditions did not differ significantly.

Similarly, we found power differences comparing the TFRs to AMWN and WN and SHAM stimulation using $TFCE_{ANOVA}$ (Supplementary Figure 2B, C, and D). These differences were prominent in two TFCE clusters, one residing in the slow wave and theta range (2 to 10.5 Hz, cluster p < 0.0001), from 0.063 to 0.883 s after stimulus onset (0 s), the other in the sigma range (11.5 to 20 Hz, cluster p = 0.0001), from 0.414 to 1.293 s after stimulus onset. Pairwise $TFCE_t$ post-hoc comparisons showed that these cluster differences were driven by the auditory stimulation conditions (Supplementary Figure 2B, and C) and that there is no significant difference between AMWN and WN (Supplementary Figure 2D).

To assess whether AMWN, WN and SHAM had differential effects on sleep spindle density we fitted a LMM to the data with fixed effects Condition (AMWN, WN, SHAM), Term (SHORT, MID) and a Condition x Term interaction (Supplementary Figure 3). The LMMs for fast and slow spindle density included random intercepts and random slopes by Term for each subject. For fast spindles, the factor Condition (F(2,52) = 5.5632, p = 0.0065) had significant effects on spindle density, while Term (F(1,13) = 4.6575, p = 0.050) and the Condition x Term interaction (F(2,52) = 0.4007, p = 0.6719) were non-significant. Post-hoc comparisons of the conditions EMMs revealed a significant decrease of fast spindles by 0.9434 ± 0.3516 spindles/30s in the AMWN condition (SHAM vs. AMWN: 0.9434 ± 0.3516 spindles/30s, $CI_{95\%}$ = [0.0954, 1.7914]; t(52 ) = 2.6833, p = 0.0259) and by 1.0750 ± 0.3516 spindles/30s in the WN condition (SHAM vs. WN: 1.0750 ± 0.3516 spindles/30s, $CI_{95\%}$ = [0.2270, 1.9231]; t(65) = 2.975, p = 0.0113) compared to SHAM, however, AMWN  did not differ significantly from WN (AMWN vs. WN: 0.1316 ± 0.3516 spindles/30s, $CI_{95\%}$ = [0.71640, 0.9796]; t(52) = 0.3744, p = 0.9258).

For slow spindle density, only Term (F(1,13) = 6.0192, p = 0.0290) was significant, while Condition (F(2,52) = 2.9574, p = 0.0608) and the Condition x Term interaction (F(2,52) = 0.4361, p = 0.6489) were unaffected.

Because sleep spindles were not differentially affected by the auditory stimulation conditions, we decided to perform the remaining analyses on pooled AMWN and WN data, referred to as STIM.

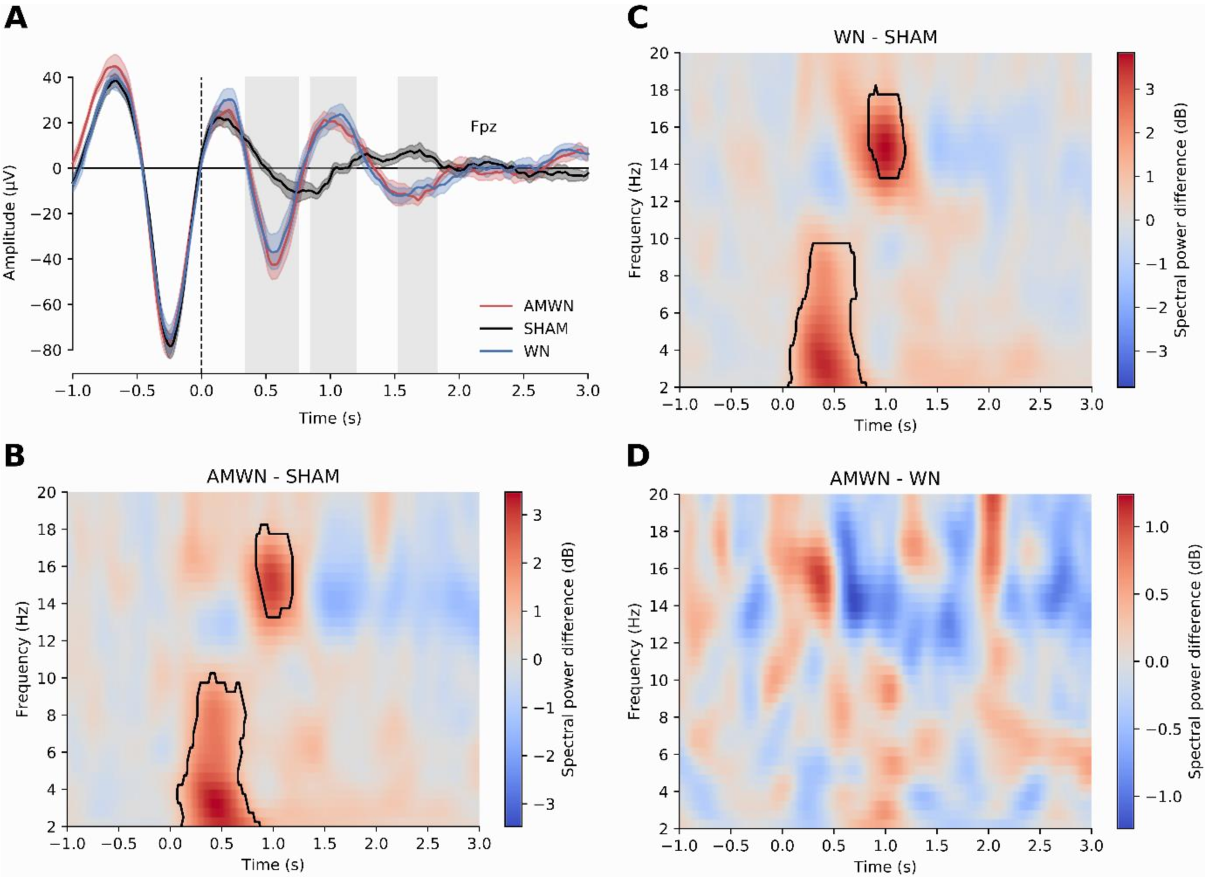


*Supplementary Figure 2 - Stimulus-locked ERPs and TFRs. (A) Plotted are grand averages of the three conditions with their respective SEMs for channel Fpz. The gray shaded areas indicate significant $TFCE_{ANOVA}$ clusters based on all channels. (B) The spectral power differences in dB between AMWN and SHAM are color-coded. Red color indicates higher power in AMWN, while blue indicates higher power in SHAM. Clusters where the spectral power differences were significantly different (obtained by $TFCE_t$) are marked by a black outline. (C) The spectral power differences in dB between WN and SHAM. Clusters where the spectral power differences were significantly different (obtained by $TFCE_t$) are marked by a black outline. (D) The spectral power differences in dB between AMWN and WN. There were no significant differences between the two stimulation modes as assessed by $TFCE_t$.*

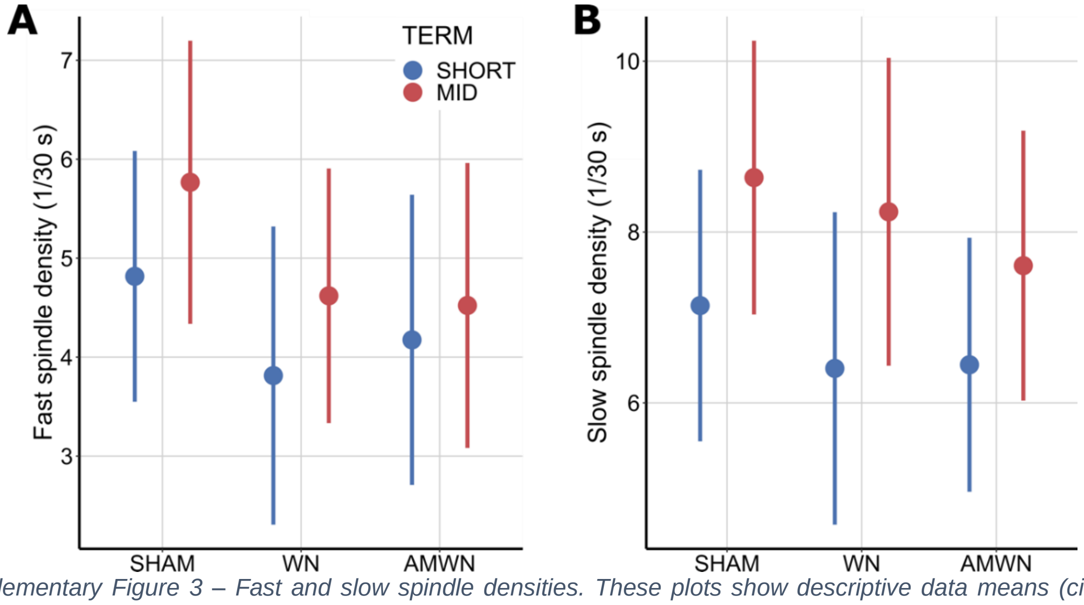


*Supplementary Figure 3 – Fast and slow spindle densities. These plots show descriptive data means (circular markers) and their respective 95% confidence intervals alongside the EMMs (diamond markers) split by Condition and Term. In the background we show the distribution of individual datapoints. (A) depicts the fast spindle density and (B) the slow spindle density.*

| Measure | Model Type | Model |
|---|---|---|
| *Fast spindle density* | LMM | FastSpindleDensity ~ Condition * Term + (Condition*Term ||SubjectID) |
| *Slow spindle density* | LMM | FastSpindleDensity ~ Condition * Term + (Term|SubjectID) |
| *Fast spindle duration* | GLMM | FastSpindleDuration ~ Condition * Term + (1|SubjectID) |
| *Slow spindle duration* | GLMM | SlowSpindleDuration ~ Condition * Term + (1|SubjectID) |
| *Fast spindle amplitude* | LMM | log(FastSpindleAmplitude) ~ Condition * Term + (1|SubjectID) |
| *Slow spindle amplitude* | LMM | log(SlowSpindleAmplitude) ~ Condition * Term + (Condition*Term ||SubjectID) |
| *Fast spindle frequency* | LMM | FastSpindleFrequency ~ Condition * Term + (Term|SubjectID) |
| *Slow spindle Frequency* | LMM | SlowSpindleFrequency ~ Condition * Term + (Condition||SubjectID) |

*Supplementary Table 1 – Statistical models fitted to fast and slow spindle measures to assess the effect of auditory stimulation (STIM vs. SHAM). This table shows the models that were fitted to fast and slow spindle measures after pooling the auditory stimulation conditions. The model structure is specified according to the afex::mixed notation (see Singmann & Kellen, 2017).*

| Measure | Model Type | Model |
|---|---|---|
| *Slow wave density* | LMM | SlowWaveDensity ~ Condition * Term + (Condition+Term |SubjectID) |
| *Slow wave duration* | LMM | log(SlowWaveDuration) ~ Condition * Term + (Term|SubjectID) |
| *Slow wave amplitude* | LMM | SlowWaveAmplitude ~ Condition * Term + (Condition||SubjectID) |

*Supplementary Table 2 - Statistical models fitted to slow wave measures to assess the effect of auditory stimulation (STIM vs. SHAM). This table shows the models that were fitted to slow wave measures after pooling the auditory stimulation conditions. The model structure is specified according to the afex::mixed notation (see Singmann & Kellen, 2017).*
